# Tunable Genotyping-By-Sequencing (tGBS®) Enables Reliable Genotyping of Heterozygous Loci

**DOI:** 10.1101/100461

**Authors:** Alina Ott, Sanzhen Liu, James C. Schnable, Cheng-Ting “Eddy” Yeh, Cassy Wang, Patrick S. Schnable

## Abstract

Most Genotyping-by-Sequencing (GBS) strategies suffer from high rates of missing data and high error rates, particularly at heterozygous sites. Tunable genotyping-by-sequencing (tGBS®), a novel genome reduction method, consists of the ligation of single-strand oligos to restriction enzyme fragments. DNA barcodes are added during PCR amplification; additional (selective) nucleotides included at the 3’-end of the PCR primers result in more genome reduction as compared to conventional GBS methods. By adjusting the number of selective bases different numbers of genomic sites can be targeted for sequencing. Because this genome reduction strategy concentrates sequencing reads on fewer sites, SNP calls are based on more reads than conventional GBS, resulting in higher SNP calling accuracy (>97-99%) even for heterozygous sites and less missing data per marker. tGBS genotyping is expected to be particularly useful for genomic selection, which requires the ability to genotype populations of individuals that are heterozygous at many loci.

## Introduction

A fundamental goal of biology is to link variation in genotype with variation in phenotype. Achieving this goal requires accurate methods for measuring both genotypes and phenotypes. The development of polymerase chain reaction (PCR) made feasible assays of genotypic variation between individuals on a scale never before achieved (Kwok 2001). The introduction of fluorescent dyes and hybridization technology have enhanced the reliability, improved the sensitivity, and increased the throughput of genotyping assays (Chee et al. 1996; Morris et al. 1996; Oliphant et al. 2002). In the last decade, advances in DNA sequencing technologies and substantial cost reduction have made it possible to assay genotypes of individual organisms via sequencing (Mardis 2011; Egan et al. 2012). Genotyping using sequence data can incorporate marker discovery and marker scoring into a single process, reducing the ascertainment bias inherent in many other PCR or hybridization-based genotyping approaches which are designed to score a pre-defined set of markers.

The most comprehensive form of genotyping using sequence data is complete resequencing of the genomes of individuals of interest at sufficient depth to identify polymorphisms. However, for most eukaryotic species this approach is still cost prohibitive. Various genome reduction strategies have been developed to target only a subset of an organism’s genome for sequencing, reducing the total amount of sequence data needed per individual. The most common genome reduction approach is to sequence genomic loci targeted by restriction enzymes (REs).

One of the first NGS-based genotyping strategies to utilize REs as a method of genome reduction was RAD-Seq (Baird et al. 2008). While RAD-Seq represented a significant advance in reducing cost and increasing throughput relative to whole genome resequencing, the initial protocol included labor intensive and costly steps such as physical shearing of DNA molecules and enzymatic end repair to process DNA. A range of protocols have subsequently been developed for employing REs as a genome reduction method, including CrOPS (van Orsouw et al. 2007), MGS (Andolfatto et al. 2011), GBS (Elshire et al. 2011), double digest RADseq (Peterson et al. 2012), 2b-RAD (Wang et al. 2012), and RESTSeq (Stolle and Moritz 2013). The majority of these innovations have been aimed at increasing the stringency of genome reduction. Even so, current methods often still target hundreds of thousands to millions of sites in a genome. As a result, given a reasonable amount of sequencing, read depth per site is often quite low, resulting in any given site remaining unsequenced in a subset of individuals, thereby resulting in high levels of missing data, low accuracy rates at heterozygous loci and reduced detection of rare alleles. Low read depth also produces higher error rates especially from heterozygous loci where smaller numbers of aligned reads increase the risk that only one of the two alleles present will be represented. This limits the use of these methods primarily to inbred lines, or requires more sequencing per individual to increase read depths, thereby reducing the advantages gained from genome reduction.

In practice, the ideal level of genome reduction will vary depending on the size of the target genome, the nature of the population being sequenced, the prevalence of polymorphic loci in this population, and the research goals. Ascertaining phylogenetic relationships can often be achieved using only a few hundred markers. Mapping QTLs within an F2 or RIL population will generally benefit from the genotyping of several thousand markers. Genome-wide association studies (GWAS) may require anywhere from tens of thousands to millions of markers depending on the level of linkage disequilibrium. In principle each of these needs could be addressed by separate genome reduction technologies. However such an approach would mean very few markers would be shared across different datasets generated for different initial aims, limiting interoperability and data reusability.

Here, we describe a new method, tunable genotyping-by-sequencing (tGBS), for genome reduction and genotyping-by-sequencing. This method provides the ability to adjust the number of targeted sites based on research goals by modifying a single primer in the protocol. In addition, unlike the genome reduction methods described above, this method removes the need for double-stranded adapters.

Our results demonstrate that sequencing reads from tGBS libraries are highly enriched at target sites and produce higher average read depths per target site given the same number of reads per sample employed by other genotyping-by-sequencing strategies. As a consequence of the high average read depth per site, a low fraction of missing data and high repeatability in SNP calls among individuals is observed, avoiding the need for extensive imputation. Finally, tGBS exhibits high accuracy in genotyping both homozygous (>97%) and heterozygous (>98%) loci, which makes genotyping-by-sequencing a more practical option in non-inbred populations such as F1BC1s and F2s widely used in both genetic research and selective breeding, including genomic selection(He et al. 2014).

## Results

### tGBS for genome reduction

During tGBS, genomic DNA is subjected to double-digestion with two enzymes in the same reaction, producing DNA fragments with a 5’ overhang on one end and a 3’ overhang on the other (Figure 1). In contrast to other methods (van Orsouw et al. 2007; Andolfatto et al. 2011; Elshire et al. 2011; Peterson et al. 2012; Wang et al. 2012; Stolle and Moritz 2013) which employ double-stranded adapters, a single-strand oligonucleotide (oligo) is ligated to each overhang. One of the oligos is unique to an individual sample and contains a DNA barcode (Qiu et al. 2003) (barcode oligo) while the other oligo is common to all samples and contains a universal sequence (universal oligo) for subsequent library construction. Following ligation, two PCR steps complete the construction of the sequencing library. For the first PCR (selective PCR), two PCR primers that partially match the ligation oligos are used. The primer matching the universal oligo (selective primer) is designed to be the reverse complement of the universal ligation oligo; however, it extends an additional 1-3 nucleotides (selective bases) at its 3’ end which can only perfectly anneal to a subset of the genomic fragments created by restriction enzyme digestion and oligo ligation, thus reducing the number of targeted sites to be amplified. As a result, genomic fragments that include the complement of the selective bases and the universal oligo will be preferentially amplified. The non-selective primer used in selective PCR matches the 5’ end of the barcode oligo. Because this primer will anneal and amplify the sequence preceding the barcode, the primer itself does not need to be designed match the barcode, reducing primer complexity and cost. For the second PCR (final PCR), two primers compatible with the appropriate sequencing platform are used to create the sequencing library.

**Figure 1.**
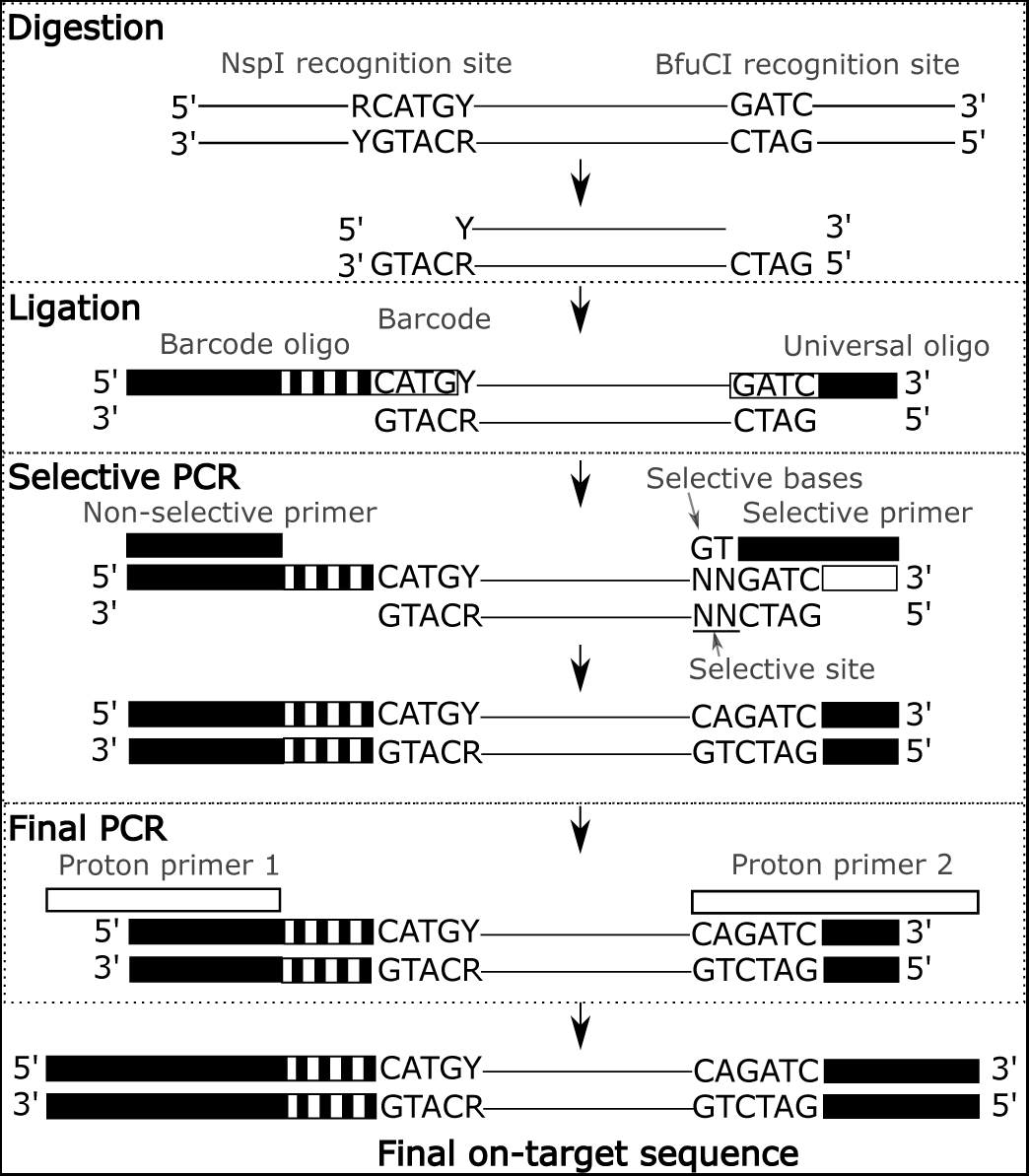
Diagram of tGBS. Digestion. Genomic DNA is digested with two restriction enzymes: NspI leaves a 3’overhang and BfuCI leaves a 5’ overhang. Ligation. Two single-strand oligos are ligated to the complementary 3’ and 5’ overhangs. The oligo matching the 3’ overhang contains a sample-specific internal barcode sequence for sample identification. The oligo matching the 5’ overhang is universal and present in every reaction for later amplification. Selective PCR. Target sites are selected using a selective primer with variable selective bases (“CA”) that match selective sites in the digested genome fragments and a non-selective primer. When properly amplified, the selective site is complementary to the selective bases. Final PCR. Primers matching the amplification primer and the selective primer which contain the full Proton adapter sequence are used for amplification of the final library. Final on-target sequence. The final sequence contains the 5’ Proton adapter sequence, an internal barcode, the NspI restriction enzyme site, the target molecule, selective bases, the BfuCI restriction enzyme site and the 3’ Proton adapter sequence. It is possible to adapt the tGBS protocol for sequencing on an Illumina instrument by redesigning the ligation oligos and PCR primers.

Based on their cutting frequencies and abilities to generate appropriate overhangs (one 5’ overhang and one 3’ overhang), the utility of NspI and BfuCI for tGBS was evaluated by simulation using the maize B73 reference genome(Schnable et al. 2009). Constraining the analysis to only non-repetitive DNA-fragments with different cut sites on each end with a total size between 100 and 300 bp yielded a total of 246,124, 44,372, and 8,645 non-repetitive DNA fragments for 1-, 2- or 3-base pairs of selective bases (T, TG, and TGT) respectively. Both the identity and number of selective bases can be adjusted to increase or decrease the expected number of fragments (Supplementary Table 1).

### Tunable genotyping-by-sequencing strongly selects for reads at target sites

The maize inbreds B73, Mo17, and the 25 parents of the Nested Association Mapping (NAM) population (Yu et al. 2008b) were genotyped via tGBS using the enzymes NspI and BfuCI and 1, 2 and 3 selective bases (Supplementary Table 2). Each level of selection is named based on the number of selective bases: e.g. genome reduction level 1 (GRL1) for 1 selective base,. An average of 6.4M (GRL1), 8.1M (GRL2), and 6.3M (GRL3) reads were generated per line. These reads were then subjected to quality trimming and aligned to the B73 reference genome.

At all GRLs, >90% of the aligned reads contain the expected restriction enzyme recognition sites. In GRL2, the selective primer had the selective bases “TG” at its 3’ end. In an ideal case, all amplified reads will be derived from sites that contain the selected “AC” sequence. However, mis-annealing of primers during PCR can lead to off-target amplification. To measure the specificity of selection during our PCR protocol, the bases at the selective site of sequenced reads were examined. Target sites in the genome contain the appropriate restriction enzyme adjacent to the selective bases (“AC” in the case of GRL2 “TG” selection), and reads that align to sites meeting these criteria are termed on-target reads.

GRL1 had the highest percent of on-target reads, with an average of 68% of the reads across all samples containing both the restriction enzyme sites and the correct selective bases based on the B73 genome. For GRL2 and GRL3 the average percent of on-target reads were 58% and 44%, respectively, across all samples (Figure 2). Note that for each additional selective base, the number of on-target sites decreases by 1/4. Therefore, even though the on-target rate was lower for GRL3 than for GRL1 and GRL2, the read depth of covered bases at on-target fragments was highest for GRL3 (Table 1). As a consequence of the size selection conducted during Proton sequencing, 68% of all uniquely aligning reads (4,248,425/6,271,577) and 88% of on-target reads (3,569,220/4,071,296) were from on-target sites between 100 and 300 bp.

**Figure 2.**
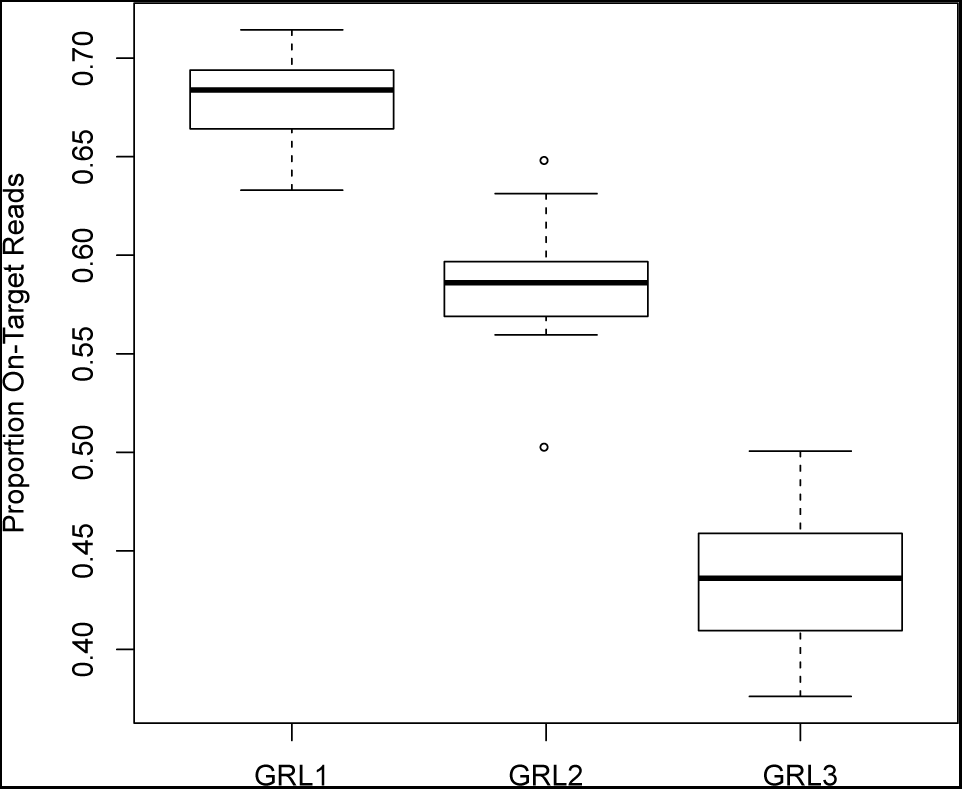
Selectivity in B73, Mo17, and the NAM founders. In the absence of selection, the proportion of random reads in the target size range from the B73 genome with “T”, “TG”, and “TGT” selection in GRL1, GLR2, and GRL3 would be 25%, 6%, and 2%, respectively.

**Table 1.** Enrichment of sequencing reads at on-target sites in B73.

| **GRL** | **% On-Target Reads** | **On-Target Interrogated Sites** | **Average Read Depth at On-Target Interrogated Sites** | **Total Reads** |
| --- | --- | --- | --- | --- |
| **1** | 65 | 68,601,482 | 10.7 | 9,109,447 |
| **2** | 65 | 16,620,747 | 37.4 | 7,903,154 |
| **3** | 44 | 5,257,786 | 113.0 | 8,428,505 |

### Application of tGBS to genotype the founders of the Nested Association Mapping (NAM) population

Genotyping diverse sets of lines is important for genome-wide association studies and genomic selection. A minimum call rate (MCR) cutoff was implemented for the 25 NAM founders. At 70% MCR, each SNP must have been genotyped in ≥ 70% of the samples. In the NAM founders, 6,665 (GRL1), 11,883 (GRL2), and 3,253 (GRL3) SNPs were identified at 70% MCR (Table 2). SNPs identified in each GRL are distributed relatively evenly across the genome (Figure 3, Supplementary Figure 2), and the number of reads per SNP site per sample had a mean of 63 and a median of 31 (Supplementary Figure 3.). The numbers of SNPs discovered in the NAM founders are not directly comparable across GRLs due to the variation in the average read number per sample (Supplementary Table 3). To overcome this limitation, a subset of NAM founders with a comparable minimum number of reads were used in the analysis described below.

**Figure 3.**
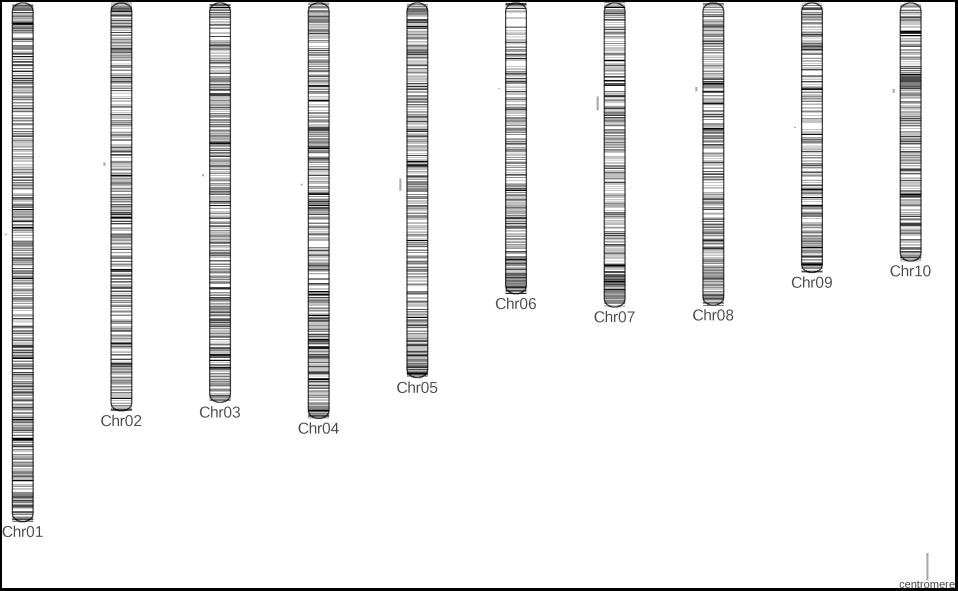
Genomic distribution of SNPs discovered in the 25 NAM founders using tGBS GRL2 at 70% MCR. Each horizontal line represents the physical position of a SNP identified by alignment to the B73 reference genome. The circles to the left of each chromosome represent the location of the centromere.

**Table 2.**
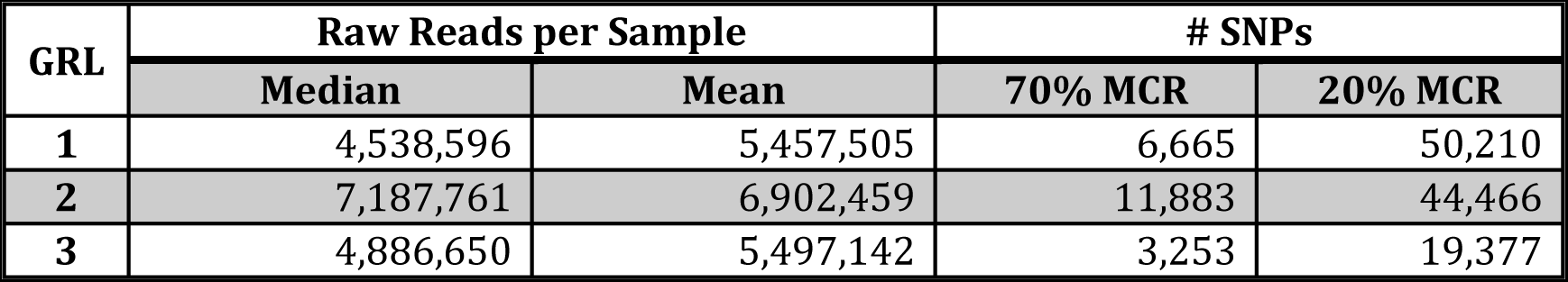
SNP identification in the 25 NAM founders.

| **GRL** | **Raw Reads per Sample** |  | **# SNPs** |  |
| --- | --- | --- | --- | --- |
|  | **Median** | **Mean** | **70% MCR** | **20% MCR** |
| **1** | 4,538,596 | 5,457,505 | 6,665 | 50,210 |
| **2** | 7,187,761 | 6,902,459 | 11,883 | 44,466 |
| **3** | 4,886,650 | 5,497,142 | 3,253 | 19,377 |

To examine the trade-offs in SNP discovery associated with reduced sequencing depth we subsampled the sequencing reads from each of the NAM founders independently. In our data set 11 of the 25 NAM founders had a sufficient number of reads across all three GRL to perform comparable subsampling (Supplementary Table 3). Therefore, only these 11 founders were included in the subsampling. From this analysis, the diminishing returns of SNP discovery with increased sequencing depth can be seen in GRL3, which begins to plateaus after 3 million raw reads. At GRL2, additional sequencing exhibits diminishing returns such that the benefits of additional sequencing begins to level off around 4 million subsampled reads (Supplementary Figure 4). GRL1 had not reached read saturation at 4 million reads (Supplementary Figure 4).

The maximum error of SNPs called in all 25 NAM founders was determined by calculating the concordance rate of tGBS SNPs with those derived from HapMap2 (Chia et al. 2012) and RNA-seq data (Yu et al. 2012) from the same lines, which were determined by whole genome resequencing and transcriptome sequencing five maize tissues for each of the NAM founders, respectively. For this analysis, individual SNPs were compared and therefore an MCR cut-off was not employed. Across the 25 founders, 90,902 GRL1, 95,028 GRL2, and 30,051 GRL3 SNPs were genotyped in all three experiments (Supplementary Table 4). To calculate minimum accuracy rates, if two of the three experiments made a concordant genotyping call at a particular location, the third non-concordant call was considered an error. tGBS had >99% concordant calls for all GRL, higher than the other two methods, supporting the accuracy of tGBS (Supplementary Table 4). Note that this approach probably overestimates genotyping errors because the lack of concordance between methods may be due to biological differences among the different pedigrees of samples used in the three experiments. Hence, the minimum SNP calling accuracy of tGBS as determined in this analysis of inbred lines is >99%.

### Genotyping recombinant inbred lines (RILs) and construction of a genetic map

The IBM RILs were developed by crossing B73 and Mo17. Random mating was performed for several generations before extensive inbreeding (Lee et al. 2002a). tGBS was conducted on 232 IBM RILs (Supplementary Table 5) using GRL2. A mean of 2.1 M reads and a median of 1.8 M reads were obtained per sample which is similar to target sequencing read numbers per SNP generally employed by other GBS protocols (Elshire et al. 2011).

The accuracy of the 70% MCR SNP calls was assessed by comparing tGBS SNP calls with Sequenom –based genotyping results (Liu et al. 2010) and RNA-seq (Li et al. 2013b) for 67 IBM RILs genotyped with all three methods, similar to the comparison performed for the NAM founders. However, unlike the NAM founders, it was possible to subdivide the genome of each RIL into segments, each of which was derived from one of the two RIL parents: B73 or Mo17. This allowed us to compare all SNPs in each of these segments to SNP calls obtained using other genotyping technologies, as opposed to just comparing individual SNP sites that were genotyped with both technologies. This approach allowed us to compare genotyping calls that were not limited by the technology with the lowest number of SNPs ( 68k in Sequenom) (Supplementary Table 6). Another difference in this analysis as compared to the analysis of the NAM founders was that heterozygosity and minor allele frequency filters (based on expected segregation patterns in RILs) were employed to exclude errors due to alignment. Following filtering, each of the three datasets was used to generate segments, which were compared to the original SNP calls used as input data for segmentation. As expected the agreement between the input data and the segmented data was high. In this analysis tGBS had a minimum accuracy of 99% (Supplementary Table 6).

Genetic maps were constructed both with and without SNP imputation at various MCR cutoffs (Figure 4). Based on Spearman rank correlation, marker orders were conserved between the genetic and physical maps (Supplementary Table 7). At 70% MCR, about 4,000 ( 90%) SNPs were mapped using both imputed and non-imputed data. As expected, SNP sets obtained using more relaxed MCR cut-offs (50% or 20% MCR) were larger. At a low MCR of 20%, imputation improved the number and percent of SNPs on the genetic map. The generation of approximately ten linkage groups corresponding to the ten maize chromosomes, the high percent of markers that are mapped, the extremely low proportion of markers assigned to an incorrect chromosome, the low estimated error rate of markers on the genetic map, and the high Spearman correlation values demonstrate that tGBS genotyping calls were quite accurate for these homozygous RILs (Supplementary Table 7).

**Figure 4.**
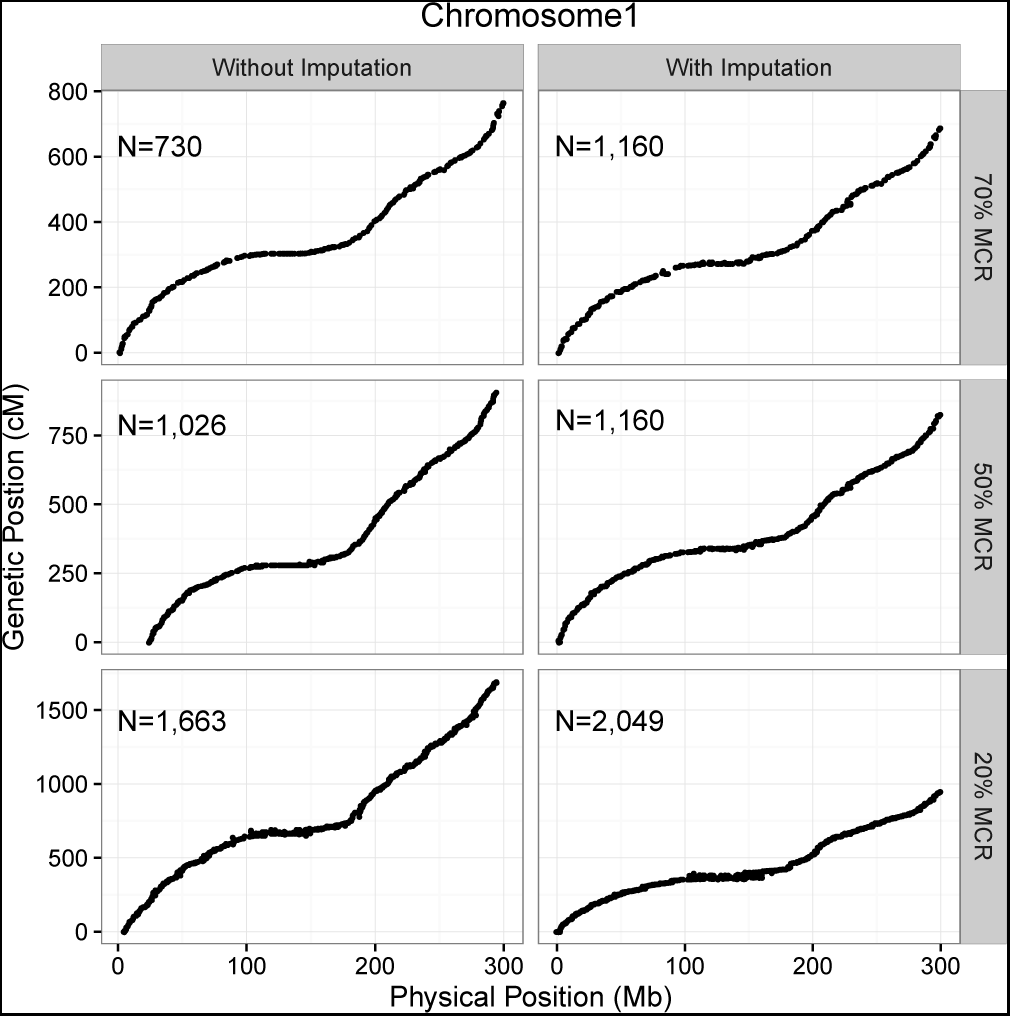
Genetic mapping in the IBM RILs. Comparisons of genetic and physical positions on chromosome 1 generated from ASMap for various MCRs, without and with LinkImpute-based imputation. Each dot represents the positon of a single SNP on a genetic and physical map.

### Application of tGBS for genotyping of heterozygous loci

To assess the accuracy of genotyping heterozygous sites, SNPs were called in 192 F2 progeny of the B73 x Mo17 cross at GRL1, GRL2, and GRL3. After filtering for MCR, minor allele frequency, and heterozygosity, the set of 70% MCR SNPs called in the F2 population were used to create of a genetic map. Similar numbers of markers (3,498/4,032, 85%), low mapping error rates (0.005), and high correlations (0.99) were obtained from the F2 data as the IBM data, indicating that tGBS performs similarly well on populations with high levels of heterozygosity (Supplementary Table 7). The presence of both homozygous and heterozygous genotypes also allowed the classification of errors identified in the F2 population as being false homozygous or false heterozygous calls using segmentation (see Methods). Only a small proportion (11,848/677,929, 1.7%) of genotyping calls at polymorphic sites were putative errors, and heterozygous calls were at least as accurate as homozygous calls (Supplementary Table 8).

Comparison of tGBS with conventional GBS To explore the advantages of tGBS relative to conventional GBS (cGBS), we compared tGBS data generated from the NAM founders presented in this paper with cGBS data generated from a large diversity panel by Romay et al. (2013)(Romay et al. 2013). Because different samples were genotyped using tGBS and cGBS, it was not possible to directly compare the SNP genotypes generated by the two technologies. Instead, for each technology we determined the number of interrogated sites and the median read depths at those sites (Methods). When controlling for library sizes, the median read depths for tGBS GRL1 and cGBS were similar. In contrast, tGBS GRL2 and GRL3 provide greater read depth per site than does cGBS (Figure 5).

**Figure 5.**
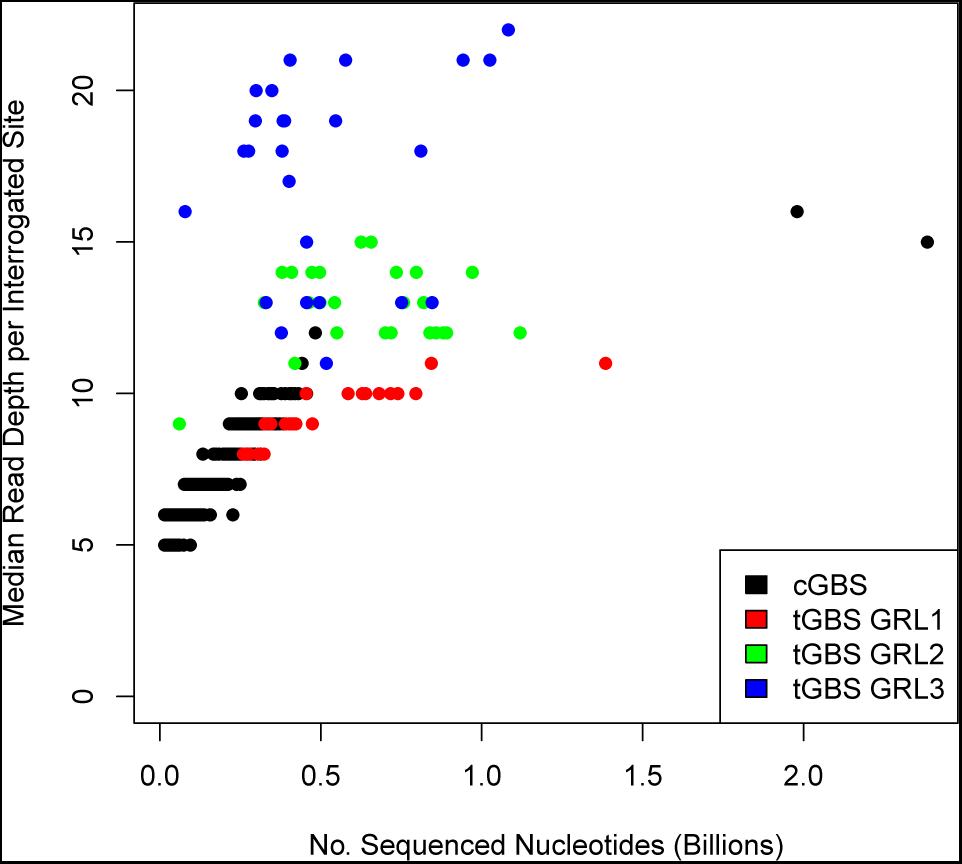
Median read depth per interrogated site for tGBS and cGBS data. Each dot represents a sample. For each GRL tGBS data were analyzed for each of 25 NAM founders. The evaluation of cGBS is based on 3,172 samples (Romay et al. 2013).

## Discussion

Here we present a novel approach to genotyping using sequence data, tGBS, which uses selection at the 3’ ends of the PCR primer to enhance genome reduction in an adjustable (tunable) manner. This method employs single-stranded oligos instead of doublestranded adaptors, which has a number of technical advantages. This genotyping approach is simple, is cost-effective, and has high accuracy at both homozygous and heterozygous sites. In addition, we have demonstrated its accuracy and reliability to genotype diversity populations, RILs, and F2s.

### Technical advantages of tGBS relative to conventional GBS

Our strategy of selecting only a subset of restriction digestion fragments for amplification and sequencing provides for flexible genome reduction. Different GRLs tune the number of target sites that will be sequenced. While fewer SNPs are obtained at higher GRL levels, the number of reads per sample necessary to saturate the genotyping of on-target SNPs is also reduced (Supplementary Figure 4, Table 1). Importantly, this results in more of the same sites across panels of samples having genotyping calls, resulting in lower levels of missing data per marker (Supplementary Figure 3, Table 2). Additionally, the increased read depth at target sites allows for accurate genotyping of both homozygous and heterozygous sites (Figure 5). This protocol could be further adapted to the Illumina TruSeq library preparation method by using DNA barcodes at both ends of amplicons in the final library, which would increase the ability to pool large numbers of samples without the need to synthesize equally large number of barcoded oligos.

### Determination of selection levels and pooling size

One of the critical decisions in any GBS experiment is how much sequencing data to generate per sample to obtain the desired number of SNPs. In maize, 12,000 and 2,000 consistently covered SNPs were obtained across 11 samples from 3 million raw GRL2 reads and 1 million raw GRL3 reads per sample, respectively (Supplementary Figure 4). In the case of the IBM RILs with GRL2 in this study, 4,293 high MCR SNPs and 10,736 low MCR SNPs were identified from an average of 2 million raw reads across all the RILs ((Supplementary Table 7). SNPs with high missing data come predominantly from off-target sites and can be imputed or disregarded, while high MCR SNPs are predominantly from on-target sites and are consistently genotyped from one experiment to the next. The appropriate GRL and number of reads per sample will vary based on the organism and project goals; however, regardless of genome complexity and diversity among individuals, sequencing depths required to on-target sites at any given threshold will be linearly related to genome size.

### Accuracy of genotyping with tGBS

Complementary methods were used to assess the accuracy of tGBS genotyping in inbreds. For the NAM founders and the IBM RILs, genotyping calls made at polymorphic sites were compared between three independent genotyping methods. Concordance between the three methods was considered an indication of accuracy while one method being non-concordant was considered an indication of an error in the non-concordant method. Requiring two out of three methods to have the same genotyping call provides a higher threshold for accuracy as the chance that an error occurs at the same site in the same sample in multiple library preparation methods is low. However, concordance as a proxy for accuracy is limited to sites that are discovered in all three methods. In addition, biological differences between samples used in the different methods can be incorrectly classified as sequencing errors. Even with error and biological differences being confounded, the accuracy estimated from the tGBS NAM concordance study was quite high at >99% (Supplementary Table 4).

While concordance in the NAM founders was limited to polymorphic sites that had been called by each of the three methods, segmentation of the IBM RILs could be used to identify regions in each RIL that are derived from either the B73 or Mo17 parent. By comparing each SNP call from multiple methods within a segment to the consensus genotype of that segment, it was possible to compare more sites. The concordance was high for all three methods, regardless of which SNP set was used to define the segments, with tGBS having a concordance >99% (Supplementary Table 6). The reported values should be considered minimum estimates of accuracy because errors and small regions with double cross overs are confounded, resulting in a potentially higher estimation of error rates. Further support for the accuracy of tGBS data is that the RIL genetic maps exhibited a high correlation with the physical marker order (>0.997), even in genetic maps constructed using unimputed SNP sets that include markers with high levels of missing data (Supplementary Table 7).

The error rates for tGBS was also found to be between 98 and 99% in a segregating F2 population using a similar segmentation-based metric (Supplementary Table 8), and the correlation between marker order on a genetic map constructed using data from the F2 individuals with the physical maize genome sequence was also >0.999 (Supplementary Table 7). The high accuracy of tGBS at heterozygous loci has the potential to increase the application of sequence-based genotyping in F2 and F1BC1 mapping populations where 50% of segregating markers are expected to be heterozygous, as well as in natural populations and obligate outcrossing species with high levels of heterozygosity. The accuracy of tGBS heterozygous genotyping will be particularly useful for conducting genomic selection, which requires the ability to genotype populations of individuals that are heterozygous at many loci.

## Materials and Methods

### Extraction of DNA Samples

DNA samples from the inbred lines B73, Mo17, and the NAM founders (Yu et al. 2008a) were extracted from 6-day old seedling tissue using the DNeasy Plant Maxi Kit [QIAGEN (Valencia, CA), No. 68163] (Supplementary Table 2). The 232 B73xMo17 recombinant inbred lines (IBM RILs) and the 192 F2 individuals(Lee et al. 2002b) were extracted from 6-day old seedling leaf tissue using the MagAttract 96 DNA Plant Core Kit [QIAGEN (Valencia, CA), No. 67163] (Supplementary Table 5). Samples were normalized using the Qubit dsDNA Broad Range Assay [ThermoFisher (Waltham, MA), no Q32853].

#### tGBS procedure

Approximately 120 ng of genomic DNA of each sample was digested with 100 units of NspI [New England Biolabs (Beverly, MA), No. R0602L] and 400 units of BfuCI [New England Biolabs (Beverly, MA), No. R0636L] in a 30 μL volume at 37°C for 1.5 hr following the manufacturer’s protocol. Unique, barcoded singlestrand oligos and a universal single-strand oligo were added to each sample for ligation with T4 DNA ligase [New England Biolabs (Beverly, MA), No. R0602L]. Ligation was performed at 16°C for 1.5 hr in a 60 μL volume following the manufacturer’s protocol. The T4 DNA ligase was inactivated by incubation at 80°C for 20 min. All digestion-ligation products were pooled and 1 mL of pooled product was purified using the QiaQuick PCR purification kit [QIAGEN (Valencia, CA), No. 28106]. The pooled, purified digestion-ligation product was used as the template for a single selective PCR reaction using a selection primer, an amplification primer, and Phusion High-Fidelity PCR Master Mix with HF Buffer [New England Biolabs (Beverly, MA), No. M0531L]. The PCR program consisted of 95°C for 3 min; 15 cycles of 98°C for 15 s, 65°C for 20 s, 72°C for 20 s; and a final extension at 72°C for 5 min. The selective PCR product was purified using Agencourt AMPure XP Beads [Beckman Coulter, Inc. (Brea, CA), No. A63880]. The purified selective PCR product was used as the template for a single, final PCR reaction using primers for the Proton platform and Phusion High-Fidelity PCR Master Mix with HF Buffer [New England Biolabs (Beverly, MA), No. M0531L]. The PCR program consisted of 98°C for 3 min; 10 cycles of 95°C for 15 s, 65°C for 20 s, 72°C for 20 s; and a final extension at 72°C for 5 min. The final PCR product was purified using Agencourt AMPure XP Beads [Beckman Coulter, Inc. (Brea, CA), No. A63880]. The purified final PCR product underwent size selection for a target of 200-300 bp using the 1.5% Agarose DNA cassette for the BluePippin [Sage Science (Beverly, MA), No. HTC2010]. The size-selected final PCR product was run on a Bioanalyzer High Sensitivity DNA chip to quantify and ensure proper size selection [Agilent Technologies (Santa Clara, CA), No. 5067-4626].

#### Debarcoding of sequencing reads and cleaning reads

Sequencing reads were analyzed with a custom Perl script which assigned each read to a sample and removed the barcode. Each debarcoded read was further trimmed to remove Proton adaptor sequences using Seqclean (sourceforge.net/projects/seqclean) and to remove potentially chimeric reads harboring internal restriction sites of NspI or BfuCI. Only reads with the correct barcodes and restriction enzyme sites were kept for further processing. These remaining reads were subjected to quality trimming. Bases with PHRED quality value <15 (out of 40) (Ewing and Green 1998; Ewing et al. 1998), i.e., error rates of ≤3%, were further removed with another custom Perl script. Each read was examined in two phases. In the first phase reads were scanned starting at each end and nucleotides with quality values lower than the threshold were removed. The remaining nucleotides were then scanned using overlapping windows of 10 bp and sequences beyond the last window with average quality value less than the specified threshold were truncated. The trimming parameters were as referred to in the trimming software, Lucy (Chou et al. 1998; Li and Chou 2004).

#### Alignment of reads to reference genome

Cleaned reads were aligned to the B73 reference genome (AGP v2) (Schnable et al. 2009) using GSNAP (Wu and Nacu 2010). Only confidently mapped reads were used for subsequent analyses, which are uniquely mapped with at least 50 bp aligned, at most 2 mismatches every 40 bp and less than a 3 bp tail for every 100 bp of read.

#### SNP discovery

The resulting confident alignments were used for SNP discovery. Reads at each potential SNP site were counted. A covered site was considered if at least five reads were counted. At each covered site, each sample was genotyped individually using the following criteria: a SNP was called as homozygous in a given sample if at least five reads supported the genotype at that site and at least 90% of all aligned reads covering that site shared the same nucleotide; a SNP was called as heterozygous in a given sample if at least two reads supported each of at least two different alleles, each of the two read types separately comprised more than 20% of the reads aligning to that site, and the sum of the reads supporting those two alleles comprised at least 90% of all reads covering the site. To compare samples with equal data, SNP discovery was performed in subsets of the data where equal numbers of randomly selected trimmed reads were processed from each sample individually.

#### Determination of selectivity

Sequencing reads obtained from Life Technology’s Proton instrument are single-end and only include the barcode, NspI digestion site, and the adjacent sequence. For this reason, selectivity could not be directly determined from reads. Selective sites for each read were predicted based on the closest BfuCI site of uniquely aligned reads in the B73 genome. On-target and off-target reads were categorized based on this selective site prediction. The number of interrogated sites was determined by identifying all the bases in the reference genome that had ≥ 5 reads uniquely aligned to that site.

In silico digestion of the B73 reference genome was performed to identify all possible NspI and BfuCI restriction enzyme fragments. Reads were aligned to this digested genome to determine which fragments have coverage.

#### Accuracy of tGBS calls

The accuracy of tGBS calls made in the NAM founders was determined by identifying concordant and disconcordant between tGBS calls and calls from TASSEL SNPs (Glaubitz et al. 2014) and RNA-sequencing SNPs (Yu et al. 2012). Polymorphic sites (i.e., at least one of the NAM founders has a non-reference allele) that were in common across the three SNP calling methods were compared. For each sample with no missing data at that site, the genotyping calls from each method were compared. If the call in one method disagreed, then the method in disagreement was considered disconcordant. Concordance was used as a proxy for accuracy.

To assess accuracy of tGBS SNP calls in the IBM RILs, tGBS SNP calls were compared to genotypes from RNA-sequencing (Liu et al. 2010) and Sequenom data in the IBM RILs (Li et al. 2013a). Because the RILs are expected to have low levels of heterozygosity and be segregating 1:1 for B73-like versus Mo17-like alleles, the tGBS and RNA-sequencing SNPs were filtered independently for sites with minor allele frequencies >0.3 and heterozygosity <0.05. A total of 67 RILs were genotyped with all three technologies and could be compared for accuracy. To increase the number of sites that could be compared between the tGBS and RNA-sequencing genotyping, segmentation was performed on each set of SNPs to identify B73-like and Mo17- like regions in each RIL. Segments were identified from each SNP set by running DNAcopy (Olshen et al. 2004) using the segment function with the parameters alpha=0.01, nperm=10000, p.method=“perm”, eta=0.01, and min.width=3. A segment genotype was determined by identifying which genotype was the majority in the given segment. The SNP genotyping calls from the each filtered SNP set were compared to the segmentation genotype from each technology.Each putative error was examined to determine the genotypes of flanking markers. If the genotype of the putative error agreed with at least one of the flanking markers, the marker was no longer considered an error. Individuals SNPs that did not match the segment genotype and had no flanking markers that would indicate the segment was generated incorrectly were considered errors.

The accuracy of tGBS calls made in the B73 x Mo17 F2 individuals was also determined by using segmentation. tGBS was performed on 192 F2 individuals at GRL2. Because an F2 population is expected to be segregating 1:2:1 at polymorphic sites with different alleles in the two parents, 4,032 SNP sites with 70% MCR, minor allele frequencies ≥ 0.35, and a proportion of heterozygous genotypes between 0.35 and 0.65 were used for segmentation. Using the same parameters for DNAcopy described above, segments of similar genotypes were identified in each of the F2 individuals Within each individual, marker genotypes that did not agree with the segment genotype (reference, heterozygous, or non-reference) were flagged as putative errors.

#### Construction of genetic maps

Genetic maps were constructed from 70% MCR, 50% MCR, and 20% MCR GRL2 SNP sets in the IBM RILs with the same filtering described for segmentation using ASMap (Taylor 2015). LinkImpute (Money et al. 2015) was run with the default settings. The imputed SNPs from LinkImpute and the unimputed SNPs for each MCR were imported into ASMap for map construction. For genetic mapping without imputation, one sample was removed due to high missing data. RILs with high similarity were detected using the comparegeno function. Four RILs were removed for having >90% similarity with another RIL. Markers with segregation distortion were identified and any markers with a p-value <1e-10 were removed. Genetic maps were constructed using the mstmap.cross function. The p-value cutoff for genetic map construction (with and without imputation) was adjusted so that 10 or more distinct linkage groups were identified (Supplementary Table 7), and the detection of bad markers was set to “yes”. The error of markers placed on the genetic map was determined by determining the maximum likelihood from a range of potential errors using R/qtl(Broman 2010).

Genetic maps were also constructed from 70% MCR GRL2 filtered SNP set for 192 B73 x Mo17 F2 individuals using ASMap. Imputation and genetic mapping were performed as described for the IBM RILs but using a more stringent p-value (<1e-5) for segregation distortion.

#### Comparison of tGBS and cGBS

cGBS data were downloaded from GenBank SRP021921 (Romay et al. 2013) Barcodes were removed and reads were trimmed and aligned to the B73 reference genome as described above for tGBS reads. For methods: Furthermore, the number of sequenced nucleotides was used to compare library size rather than number of reads as the cGBS data was generated using Illumina while tGBS data was generated using Proton.

#### Data availability

The sequencing data generated in this study are available in the Sequence Read Archive with the identifiers SRP095743 (RILs), SRP095751, SRP095750, SRP095749 (NAM GRL1, GRL2, and GRL3 respectively), and SRP095555 (F2s).

## Acknowledgments

The authors would like to thank Molly Parsons and Samantha Hoesel for assistance with wet lab experiments. Alina Ott was supported in part by fellowships from the Office of Biotechnology, Iowa State University and the National Science Foundation Graduate Research Fellowship (Grant No. DGE1247194). Any opinions, findings, and conclusions or recommendations expressed in this material are those of the authors(s) and do not necessarily reflect the views of the National Science Foundation. Finally, we would like to dedicate this paper to the memory of Matt Hickenbotham from Thermo Fisher who passed away in June 2015 and enthusiastically supported the development of this technology.

## Author Contributions

S.L and P.S.S designed the project. A.O. and S.L. conducted the experiments. A.O., S.L., and C.-T.Y. analyzed the data. A.O., S.L.,. J.C.S., C.W and P.S.S. interpreted the results. A.O., S.L., J.C.S. and P.S.S. wrote the manuscript.

## Supplemental Figures and Tables

**Supplementary Figure 1.**
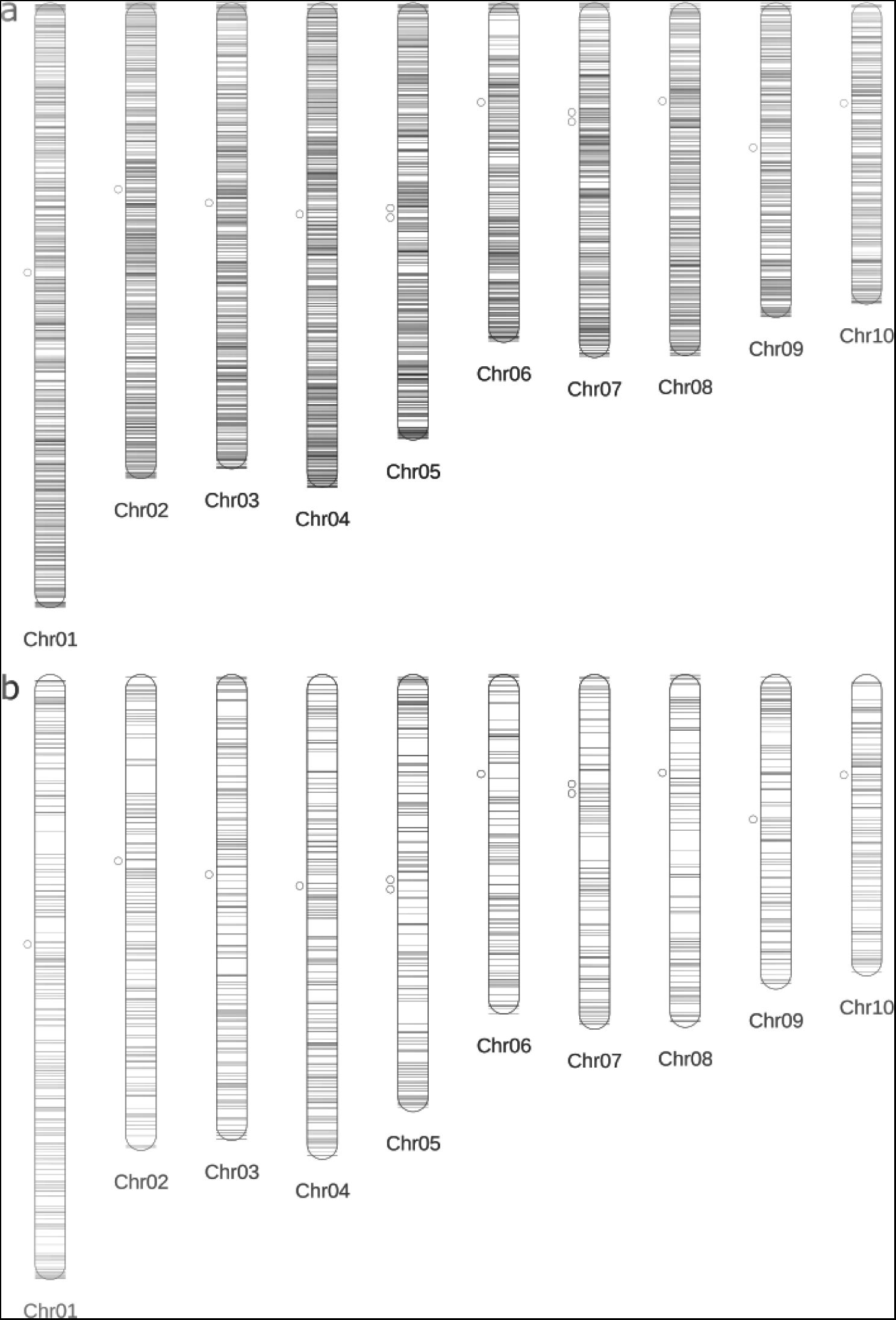
Locations of tGBS SNPs discovered in the 25 NAM founders from a.) GRL1 and b.) GRL3 at 70% MCR with each horizontal line representing the physical position of a SNP identified via alignment to the B73 reference genome.

**Supplementary Figure 2.**
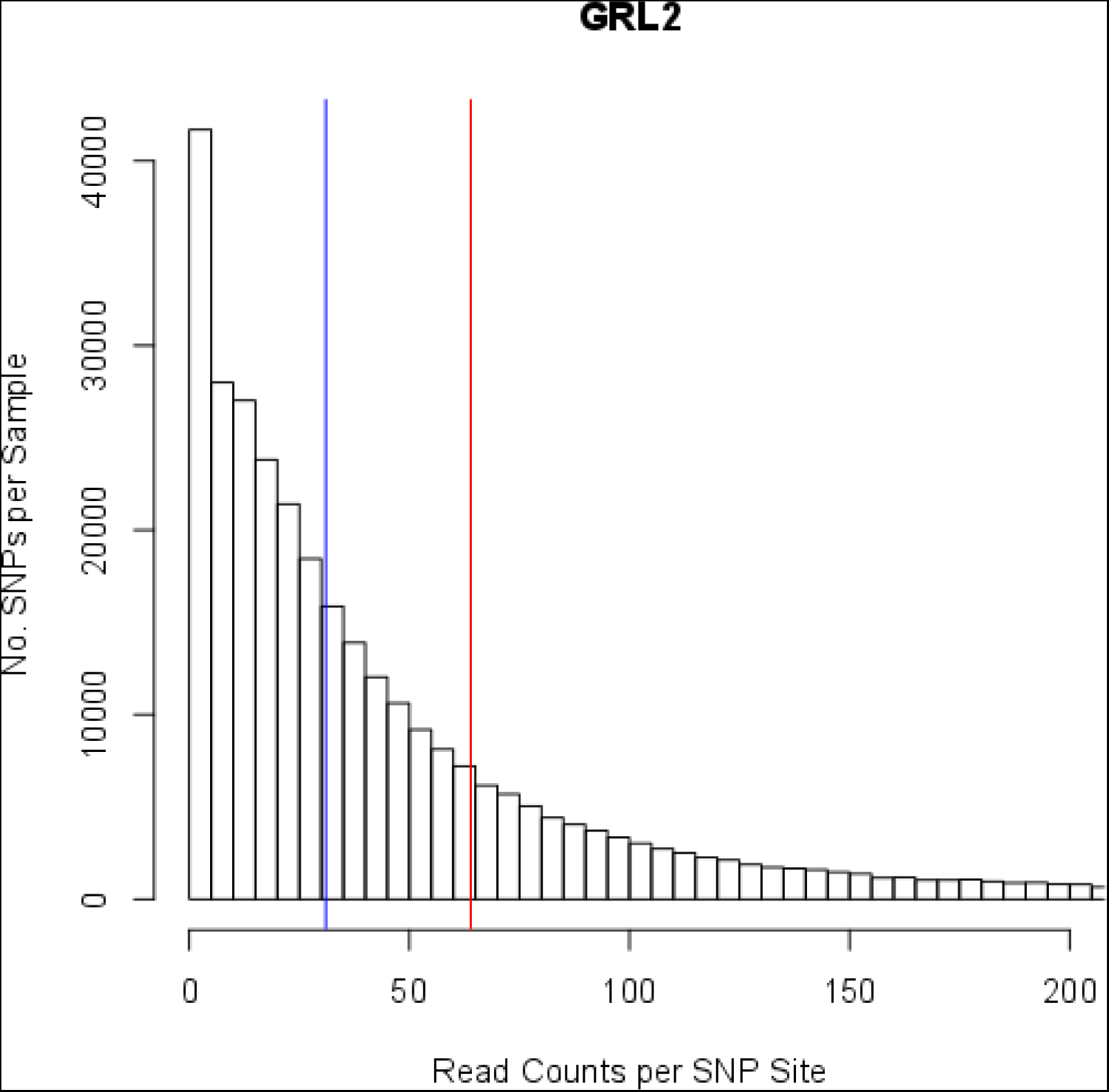
Read counts per SNP site per sample from the NAM MCR 70 SNP set. SNPs with > 200 reads per site are truncated. The mean (red line) and median (blue line) reads per site are based on all SNPs.

**Supplementary Figure 3.**
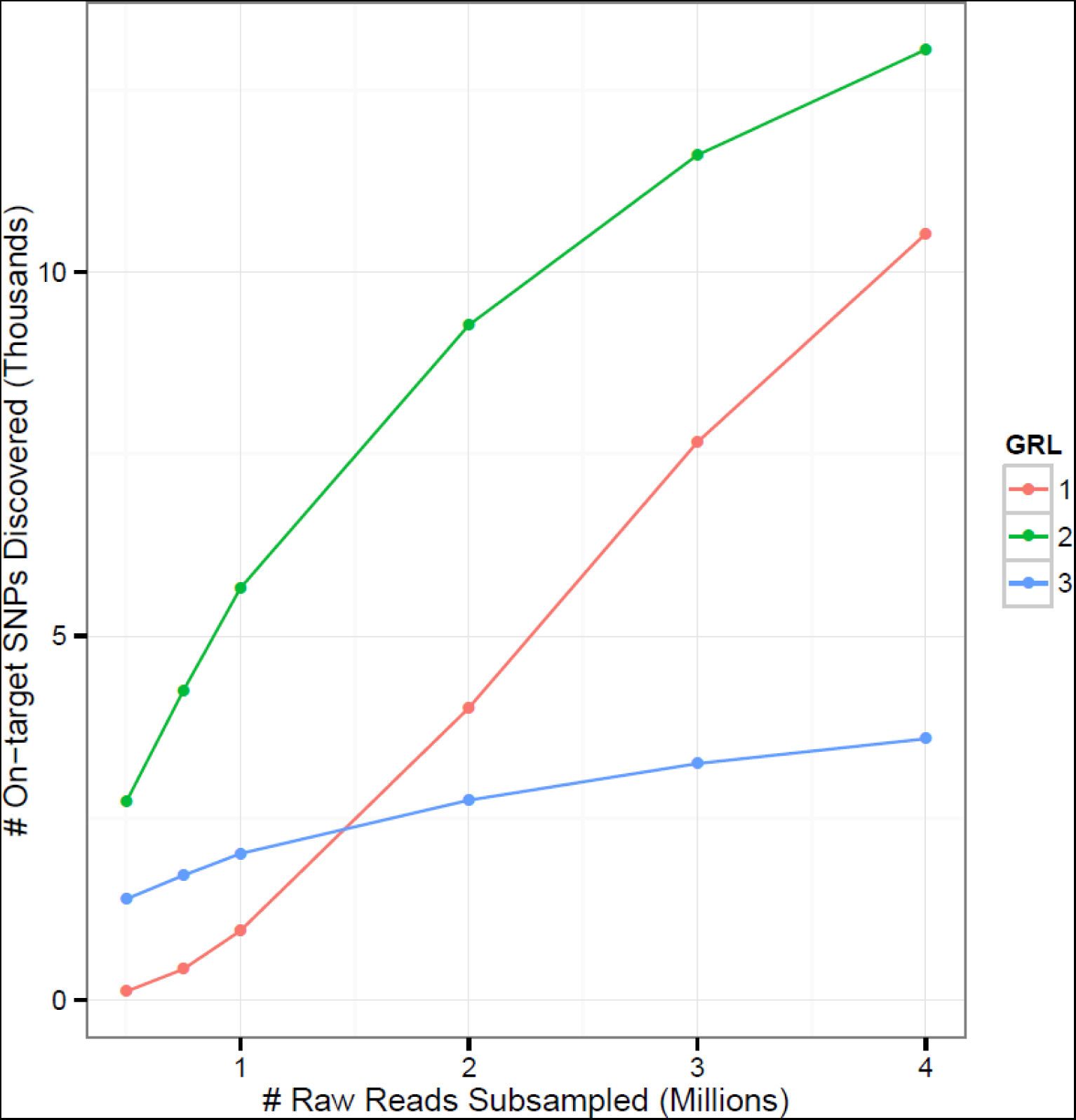
SNP discovery (70% MCR) from 11 NAM founders with varying numbers of subsampled sequenced reads.

**Supplementary Table 1.**
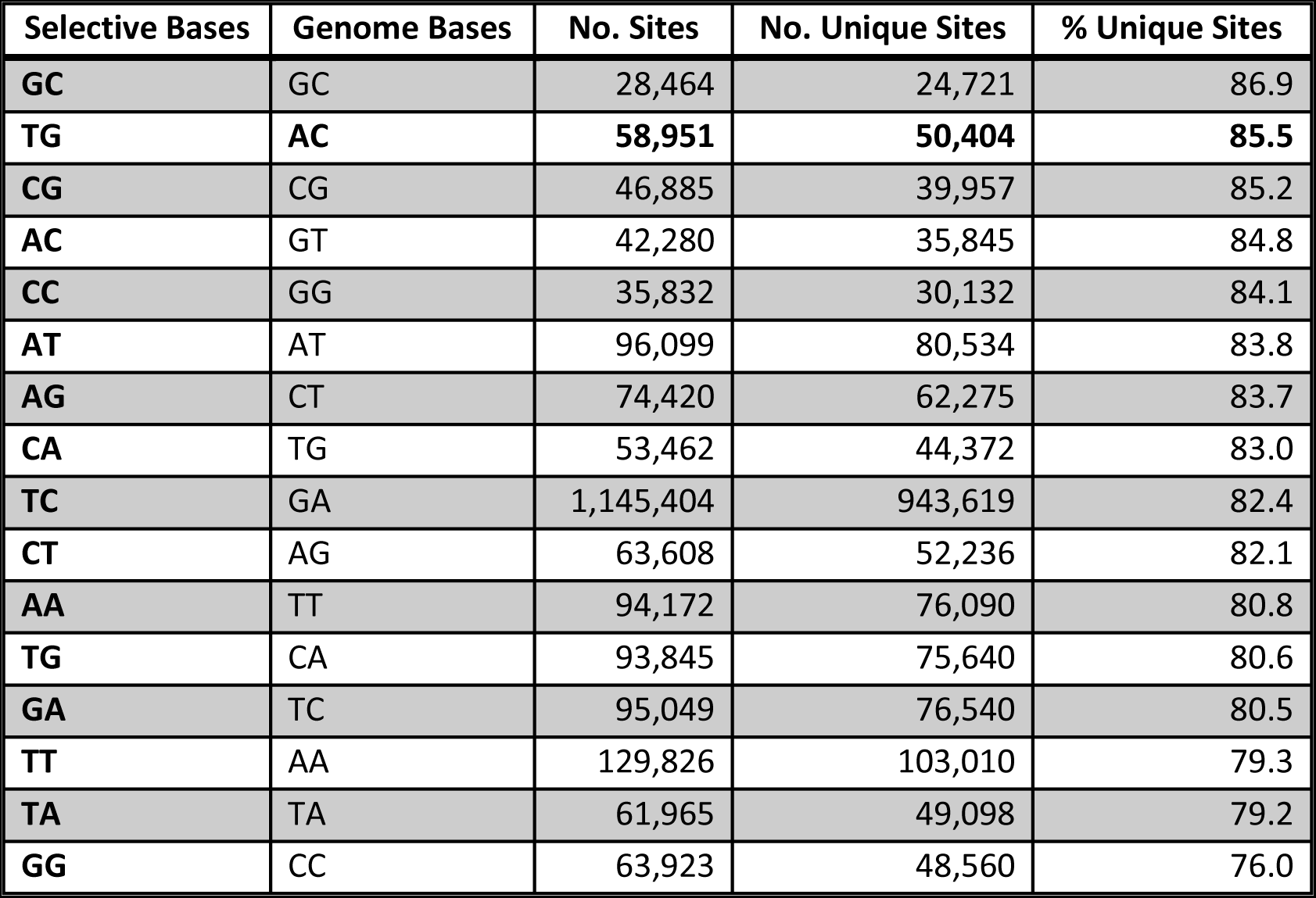
In silico digestion of the maize genome for all two-base combinations.

| Selective Bases | Genome Bases | No. Sites | No. Unique Sites | % Unique Sites |
| --- | --- | --- | --- | --- |
| GC | GC | 28,464 | 24,721 | 86.9 |
| TG | AC | **58,951** | **50,404** | **85.5** |
| CG | CG | 46,885 | 39,957 | 85.2 |
| AC | GT | 42,280 | 35,845 | 84.8 |
| CC | GG | 35,832 | 30,132 | 84.1 |
| AT | AT | 96,099 | 80,534 | 83.8 |
| AG | CT | 74,420 | 62,275 | 83.7 |
| CA | TG | 53,462 | 44,372 | 83.0 |
| TC | GA | 1,145,404 | 943,619 | 82.4 |
| CT | AG | 63,608 | 52,236 | 82.1 |
| AA | TT | 94,172 | 76,090 | 80.8 |
| TG | CA | 93,845 | 75,640 | 80.6 |
| GA | TC | 95,049 | 76,540 | 80.5 |
| TT | AA | 129,826 | 103,010 | 79.3 |
| TA | TA | 61,965 | 49,098 | 79.2 |
| GG | CC | 63,923 | 48,560 | 76.0 |

**Supplementary Table 2.** Pedigrees of NAM, B73, and Mo17.

| Genotype | Pedigree<sup>1</sup> |  |  |
| --- | --- | --- | --- |
|  | GRL 1 | GRL 2 | GRL 3 |
| B97 | 08NAM-1150-21 | 08NAM-1150-21 | 08NAM-1150-21 |
| CML103 | 07-1589-8 | 07-1589-8 | 07-1589-8 |
| CML228 | 14-2879-3 | 14-2879-3 | 14-2879-3 |
| CML247 | 11-6199-21 | 11-6199-21 | 11-6199-21 |
| CML277 | 10g-1032-3 | 10g-1032-3 | 10g-1032-3 |
| CML322 | 10g-1090-1 | 10g-1090-1 | 10g-1090-1 |
| CML333 | 10g-1091-1 | 10g-1091-1 | 10g-1091-1 |
| CML52 | 10g-1092-6 | 10g-1092-6 | 10g-1092-6 |
| CML69 | 07g-1134-8 | 07g-1134-8 | 11-6204-22 |
| Hp301 | 09-4244-2 | 09-4244-2 | 09-4244-2 |
| IL14H | 09-4245-2 | 09-4245-2 | 09-4245-2 |
| Ki11 | 07g-1138-10 | 07g-1138-10 | 07g-1138-10 |
| Ki3 | 07g-1139-4 | 07g-1139-4 | 07g-1139-4 |
| Ky21 | 14-2890-5 | 14-2890-5 | 14-2890-5 |
| M162W | 14-2891-5 | 14-2891-5 | 14-2891-5 |
| M37W | 09-4247-1 | 09-4247-1 | 09-4247-1 |
| Mo18W | 13B-6091 | 13B-6091 | 13B-6091 |
| MS71 | 14B-1411 | 14B-1411 | 14B-1411 |
| NC350 | Ac 3700 | Ac 3700 | Ac 3700 |
| NC358 | 07-1600-11 | 07-1600-11 | 07-1600-11 |
| Oh43 | 08NAM-1170-21 | 08NAM-1170-21 | 08NAM-1170-21 |
| Oh7B | 14-2875-5 | 14-2875-5 | 14-2875-5 |
| P39 | 08NAM-1172-21 | 08NAM-1172-21 | 08NAM-1172-21 |
| Tx303 | 10g-1034-1 | 10g-1034-1 | 10g-1034-1 |
| Tzi8 | 13B-165-2 | 13B-165-2 | 13B-165-1 |
| B73 | 08-3868-3 | 08-3868-3 | 08-3868-3 |
| Mo17 | 08-3877-2 | 08-3877-2 | 08-3877-2 |

<sup>1</sup> Inbred line pedigrees maintained by selfing.

**Supplementary Table 3.**
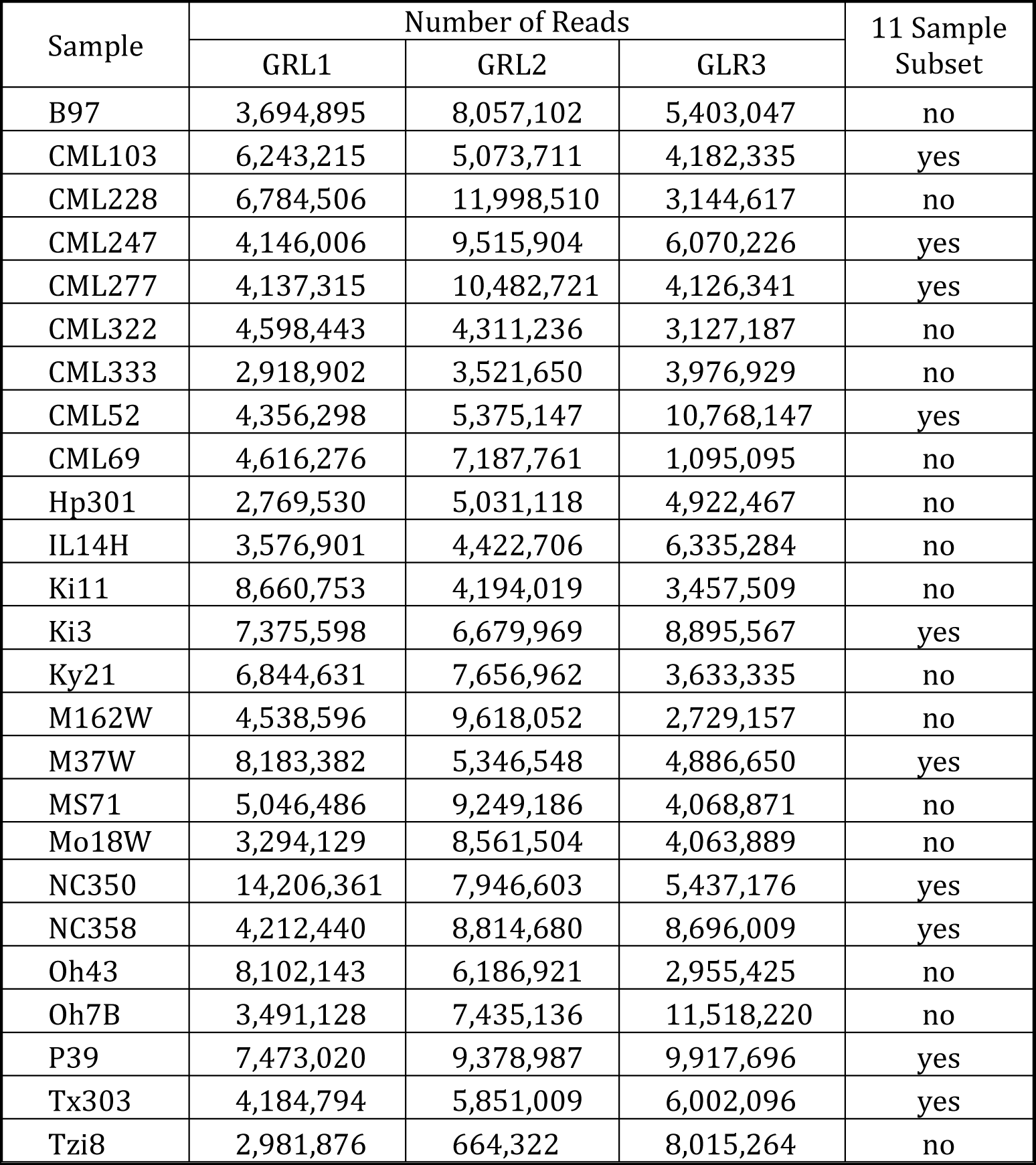
NAM founder reads per sample for GRL1-3. Founders with > 4M reads for all three GRL were used in the 11 sample subset.

**Supplementary Table 4.** Concordant SNP calls summed across the NAM founders.

| SNP Source | Concordant SNPs (%) |  |  |
| --- | --- | --- | --- |
|  | GRL1<sup>1</sup> | GRL2<sup>2</sup> | GRL3<sup>3</sup> |
| tGBS | 90,790 (99.88%) | 94,856 (99.82%) | 29,959 (99.69 %) |
| RNA-Seq | 89,215 (98.42%) | 93,537 (98.43%) | 29,598 (98.49%) |
| HapMap2 | 89,452 (98.40%) | 93,383 (98.27%) | 29,512 (98.21%) |

<sup>1</sup> 90,902 total SNPs in GRL1.
<sup>2</sup> 95,028 total SNPs in GRL2.
<sup>3</sup> 30,051 total SNPs in GRL3.

**Supplementary Table 5.** Pedigrees of IBM RILs.

| **Tissue Collection** | **Pedigree<sup>1</sup>** | **Genotype** |
| --- | --- | --- |
| 14B-1572 | 05-6152-1 | M0001 |
| 14B-1574 | 05-6165-4 | M0004 |
| 14B-1575 | 05-6304-2 | M0005 |
| 14B-1576 | 05-6166-1 | M0006 |
| 14B-1577 | 05-6155-1 | M0007 |
| 14B-1578 | 05-6317-1 | M0008 |
| 14B-1651 | 05-6306-2 | M0010 |
| 14B-1652 | 05-6318-4 | M0011 |
| 14B-1579 | 05-6157-1 | M0012 |
| 14B-1654 | 05-6308-3 | M0014 |
| 14B-1580 | 05-6159-1 | M0016 |
| 14B-1581 | 05-6321-1 | M0017 |
| 14B-1582 | 05-6322-1 | M0022 |
| 14B-1584 | 05-6323-1 | M0024 |
| 14B-1586 | 05-6324-1 | M0026 |
| 14B-1587 | 05-6187-1 | M0028 |
| 14B-1589 | 05-6177-1 | M0031 |
| 14B-1590 | 05-6189-1 | M0032 |
| 14B-1591 | 05-6178-6 | M0033 |
| 14B-1658 | 05-6340-2 | M0034 |
| 14B-1659 | 05-6191-3 | M0037 |
| 14B-1594 | 05-6192-2 | M0040 |
| 14B-1660 | 05-6181-3 | M0042 |
| 14B-1661 | 05-6193-5 | M0043 |
| 14B-1595 | 05-6182-2 | M0044 |
| 14B-1662 | 05-6194-4 | M0045 |
| 14B-1596 | 05-6183-1 | M0046 |
| 14B-1663 | 05-6195-5 | M0047 |
| 14B-1597 | 05-6184-3 | M0048 |
| 14B-1598 | 05-6196-1 | M0051 |
| 14B-1599 | 05-6185-5 | M0052 |
| 14B-1665 | 05-6198-2 | M0055 |
| 14B-1666 | 05-6351-1 | M0056 |
| 14B-1667 | 05-6364-1 | M0059 |
| 14B-1871 | 05-6353-1 | M0060 |
| 14B-1603 | 05-6365-1 | M0061 |
| 14B-1668 | 05-6204-3 | M0064 |
| 14B-1605 | 05-6367-2 | M0067 |

<sup>1</sup> RIL pedigrees maintained by selfing.

|  |  |  |
| --- | --- | --- |
| 14B-1670 | 05-6206-3 | M0069 |
| 14B-1673 | 05-6219-3 | M0073 |
| 14B-1606 | 05-6208-1 | M0075 |
| 14B-1607 | 05-6209-1 | M0077 |
| 14B-1674 | 05-6221-2 | M0078 |
| 14B-1608 | 05-6210-1 | M0079 |
| 14B-1675 | 05-6372-1 | M0080 |
| 14B-1609 | 05-6211-1 | M0081 |
| 14B-1610 | 05-6223-3 | M0083 |
| 14B-1676 | 05-6361-3 | M0084 |
| 14B-1677 | 05-6374-1 | M0085 |
| 14B-1678 | 05-6375-2 | M0086 |
| 14B-1679 | 05-6387-4 | M0088 |
| 14B-1680 | 05-6226-2 | M0090 |
| 14B-1681 | 05-6238-2 | M0091 |
| 14B-1682 | 05-6377-3 | M0092 |
| 14B-1683 | 05-6239-4 | M0093 |
| 14B-1684 | 05g-1264-4 | M0095 |
| 14B-1686 | 05-6379-2 | M0097 |
| 14B-1687 | 05-6241-1 | M0098 |
| 14B-1611 | 05-6230-4 | M0099 |
| 14B-1688 | 05-6242-4 | M0100 |
| 14B-1691 | 06-2367-3 | M0103 |
| 14B-1692 | 05-6244-1 | M0104 |
| 14B-1693 | 05-6383-1 | M0105 |
| 14B-1694 | 05-6245-1 | M0106 |
| 14B-1695 | 05-6384-4 | M0107 |
| 14B-1699 | 05-6386-2 | M0114 |
| 14B-1612 | 05-6402-4 | M0116 |
| 14B-1702 | 05-6253-1 | M0118 |
| 14B-1704 | 05-6254-3 | M0121 |
| 14B-1705 | 05-6266-4 | M0122 |
| 14B-1707 | 05-6267-5 | M0124 |
| 14B-1708 | 05-6256-7 | M0125 |
| 14B-1709 | 05-6418-5 | M0126 |
| 14B-1710 | 05-6257-1 | M0127 |
| 14B-1711 | 05-6269-1 | M0129 |
| 14B-1712 | 05-6408-2 | M0130 |
| 14B-1713 | 07-1721-6 | M0131 |
| 14B-1714 | 05-6409-2 | M0132 |
| 14B-1715 | 05-6271-1 | M0133 |
| 14B-1716 | 05-6410-3 | M0138 |
| 14B-1717 | 05-6272-1 | M0139 |
| 14B-1719 | 05-6273-5 | M0142 |
| 14B-1720 | 05-6412-2 | M0143 |
| 14B-1721 | 05-6274-2 | M0144 |
| 14B-1722 | 05g-1266-3 | M0145 |
| 14B-1723 | 05-6437-3 | M0147 |
| 14B-1724 | 05-6276-3 | M0149 |
| 14B-1725 | 06-2369-1 | M0150 |
| 14B-1728 | 05-6278-3 | M0154 |
| 14B-1729 | 05-6440-1 | M0155 |
| 14B-1730 | 05-6279-4 | M0156 |
| 14B-1731 | 05-6291-2 | M0157 |
| 14B-1732 | 05-6280-1 | M0159 |
| 14B-1734 | 05-6443-2 | M0161 |
| 14B-1736 | 05-6433-3 | M0163 |
| 14B-1613 | 05-6295-1 | M0165 |
| 14B-1737 | 05-6284-3 | M0166 |
| 14B-1739 | 05-6435-4 | M0168 |
| 14B-1740 | 05-6447-1 | M0169 |
| 14B-1741 | 05-6448-1 | M0172 |
| 14B-1743 | 05-6451-2 | M0173 |
| 14B-1744 | 05-6463-1 | M0174 |
| 14B-1745 | 05-6602-1 | M0176 |
| 14B-1746 | 05-6464-3 | M0177 |
| 14B-1747 | 05-6603-3 | M0178 |
| 14B-1748 | 05-6465-2 | M0179 |
| 14B-1749 | 05-6604-3 | M0180 |
| 14B-1750 | 05-6466-5 | M0181 |
| 14B-1752 | 05-6467-1 | M0187 |
| 14B-1753 | 07-1725-7 | M0188 |
| 14B-1754 | 05-6618-2 | M0189 |
| 14B-1755 | 05-6607-2 | M0191 |
| 14B-1756 | 05-6469-3 | M0192 |
| 14B-1757 | 05-6458-3 | M0194 |
| 14B-1759 | 05-6609-3 | M0196 |
| 14B-1760 | 05-6471-1 | M0197 |
| 14B-1761 | 05-6460-1 | M0198 |
| 14B-1762 | 05-6472-2 | M0199 |
| 14B-1764 | 05-6473-1 | M0201 |
| 14B-1765 | 05-6462-1 | M0203 |
| 14B-1766 | 05-6624-1 | M0204 |
| 14B-1768 | 05-6487-4 | M0206 |
| 14B-1769 | 06-2388-4 | M0208 |
| 14B-1770 | 05-6488-1 | M0209 |
| 14B-1772 | 05-6489-3 | M0212 |
| 14B-1773 | 05-6628-3 | M0213 |
| 14B-1774 | 05-6640-3 | M0214 |
| 14B-1775 | 05-6629-3 | M0215 |
| 14B-1776 | 05-6491-1 | M0216 |
| 14B-1777 | 05-6480-3 | M0217 |
| 14B-1778 | 05-6492-2 | M0218 |
| 14B-1779 | 05-6631-1 | M0219 |
| 14B-1780 | 05-6493-2 | M0220 |
| 14B-1781 | 05-6632-3 | M0221 |
| 14B-1782 | 05-6644-1 | M0222 |
| 14B-1783 | 05-6633-2 | M0223 |
| 14B-1784 | 05g-1267-2 | M0225 |
| 14B-1785 | 05-6484-2 | M0228 |
| 14B-1786 | 05-6646-1 | M0229 |
| 14B-1787 | 05-6485-5 | M0230 |
| 14B-1788 | 05-6497-1 | M0231 |
| 14B-1789 | 05-6648-3 | M0233 |
| 14B-1790 | 05-6502-3 | M0234 |
| 14B-1791 | 05-6664-1 | M0235 |
| 14B-1792 | 05-6653-2 | M0236 |
| 14B-1793 | 05-6515-3 | M0237 |
| 14B-1794 | 05-6654-1 | M0238 |
| 14B-1795 | 07-1726-4 | M0239 |
| 14B-1796 | 05-6505-3 | M0240 |
| 14B-1797 | 05-6667-2 | M0241 |
| 14B-1799 | 05-6518-3 | M0245 |
| 14B-1616 | 05-6507-3 | M0248 |
| 14B-1800 | 05-6669-2 | M0249 |
| 14B-1801 | 05-6658-3 | M0251 |
| 14B-1802 | 05-6670-2 | M0252 |
| 14B-1804 | 05-6671-1 | M0256 |
| 14B-1805 | 05-6521-3 | M0258 |
| 14B-1806 | 05-6511-4 | M0259 |
| 14B-1807 | 05-6673-1 | M0260 |
| 14B-1617 | 05-6662-3 | M0262 |
| 14B-1808 | 05-6674-1 | M0263 |
| 14B-1618 | 05-6525-1 | M0264 |
| 14B-1620 | 05-6526-1 | M0266 |
| 14B-1621 | 05-6688-2 | M0267 |
| 14B-1622 | 05-6774-3 | M0269 |
| 14B-1809 | 05-6677-1 | M0270 |
| 14B-1810 | 05-6689-3 | M0271 |
| 14B-1811 | 05-6528-2 | M0272 |
| 14B-1812 | 05-6540-2 | M0273 |
| 14B-1813 | 05-6529-2 | M0274 |
| 14B-1814 | 05-6541-2 | M0275 |
| 14B-1815 | 05-6542-2 | M0277 |
| 14B-1816 | 05-6681-2 | M0279 |
| 14B-1818 | 05-6694-4 | M0282 |
| 14B-1625 | 05-6803-2 | M0284 |
| 14B-1819 | 05-6802-2 | M0285 |
| 14B-1626 | 05-6545-1 | M0287 |
| 14B-1628 | 05-6546-4 | M0289 |
| 14B-1821 | 06-2391-2 | M0290 |
| 14B-1822 | 05-6697-1 | M0291 |
| 14B-1823 | 06-2392-5 | M0292 |
| 14B-1824 | 05-6548-2 | M0293 |
| 14B-1825 | 05-6701-1 | M0294 |
| 14B-1826 | 05-6713-1 | M0295 |
| 14B-1630 | 05-6553-2 | M0298 |
| 14B-1828 | 05-6554-2 | M0300 |
| 14B-1829 | 05-6716-1 | M0302 |
| 14B-1631 | 06-2375-6 | M0303 |
| 14B-1830 | 05-6567-2 | M0304 |
| 14B-1831 | 05-6706-2 | M0305 |
| 14B-1832 | 05-6568-3 | M0306 |
| 14B-1833 | 05-6557-2 | M0307 |
| 14B-1834 | 05-6719-2 | M0308 |
| 14B-1835 | 05-6558-3 | M0309 |
| 14B-1632 | 05-6570-1 | M0310 |
| 14B-1633 | 05-6559-5 | M0311 |
| 14B-1836 | 05-6721-4 | M0312 |
| 14B-1838 | 05-6710-1 | M0313 |
| 14B-1839 | 05-6561-4 | M0317 |
| 14B-1841 | 05-6562-3 | M0320 |
| 14B-1635 | 05-6575-1 | M0322 |
| 14B-1636 | 05-6587-1 | M0323 |
| 14B-1843 | 05-6726-2 | M0324 |
| 14B-1638 | 05-6577-7 | M0326 |
| 14B-1639 | 05-6578-1 | M0328 |
| 14B-1845 | 05-6590-1 | M0330 |
| 14B-1846 | 05-6729-5 | M0331 |
| 14B-1847 | 05-6591-1 | M0334 |
| 14B-1848 | 05-6730-1 | M0335 |
| 14B-1640 | 05-6592-1 | M0337 |
| 14B-1849 | 05-6731-2 | M0338 |
| 14B-1850 | 05-6743-4 | M0339 |
| 14B-1641 | 05-6594-3 | M0341 |
| 14B-1851 | 05-6804-3 | M0342 |
| 14B-1854 | 05-6565-1 | M0348 |
| 14B-1855 | 05-6746-2 | M0350 |
| 14B-1856 | 05g-1268-2 | M0351 |
| 14B-1642 | 05-6597-1 | M0352 |
| 14B-1858 | 05g-1269-2 | M0354 |
| 14B-1859 | 05-6752-5 | M0355 |
| 14B-1860 | 05-6788-1 | M0357 |
| 14B-1861 | 05g-1270-2 | M0358 |
| 14B-1862 | 05-6754-2 | M0362 |
| 14B-1863 | 05-6790-2 | M0364 |
| 14B-1643 | 05-6779-1 | M0365 |
| 14B-1645 | 05-6792-7 | M0369 |
| 14B-1865 | 05-6793-3 | M0375 |
| 14B-1866 | 05-6758-1 | M0376 |
| 14B-1867 | 07g-1151-1 | M0377 |
| 14B-1646 | 05-6759-1 | M0378 |
| 14B-1648 | 05-6761-2 | M0382 |
| 14B-1649 | 05-6797-3 | M0383 |

**Supplementary Table 6.** Percent agreement among SNP calls generated via tGBS, RNA-Seq, and Sequenom for the 67 IBM RILs that were genotyped with all three technologies. Concordance between input SNP calls derived from a given genotyping technology and SNP calls derived following segmentation of the same input data are shaded in gray. Non-shaded cells show the concordance between input SNPs and SNP calls derived following segmentation of input SNPs generated with one of the other two genotyping technologies.

| No. SNPs in Agreement<br>(% Agreement) |  | Segmented SNP Calls |  |  |  |
| --- | --- | --- | --- | --- | --- |
|  |  | tGBS | Sequenom | RNA-Seq | Total SNPs |
| Input<br>SNP Calls | tGBS | 246,344 (99.48) | 246,515 (99.55) | 246,595 (98.58) | 247,628 |
|  | Sequenom | 68,107 (99.93) | 68,079 (99.89) | 68,122 (99.95) | 68,154 |
|  | RNA-Seq | 9,284,580 (99.68) | 9,284,297 (99.68) | 9,279,703 (99.69) | 9,314,537 |

**Supplementary Table 7.** Summary of genetic maps constructed using ASMap.

| Pop. | MCR | Imp. | No. Markers |  |  |  |  | Map P-Value<sup>6</sup> | No. LG | Map Size (cM) | Average Marker Spacing (cM) | Mapped Genotyping Error Rate of Markers | Spearman Rank Correlation of Physical and Genetic Marker Order |
| --- | --- | --- | --- | --- | --- | --- | --- | --- | --- | --- | --- | --- | --- |
|  |  |  | Input | Filtered for Segregation Distortion | Mapped (%) | Incorrect LG | Un-mapped |  |  |  |  |  |  |
| IBM | 70 | no | 4,293 | 0 | 3,942 (91.9) | 9 | 351 | 1E-12 | 11 | 3506 | 0.9 | 0.0005 | 0.9982 |
| IBM | 70 | yes | 4,293 | 55 | 3,856 (87.7) | 9 | 437 | 1E-10 | 10 | 2525 | 0.7 | 0.0010 | 0.9999 |
| IBM | 50 | no | 6,696 | 0 | 6,010 (89.8) | 14 | 686 | 1E-20 | 10 | 5007 | 0.8 | 0.0010 | 0.9999 |
| IBM | 50 | yes | 6,696 | 57 | 6,200 (92.6) | 14 | 439 | 1E-14 | 10 | 3210 | 0.5 | 0.0010 | 0.9999 |
| IBM | 20 | no | 10,736 | 0 | 8,842 (82.4) | 23 | 1,894 | 1E-27 | 12 | 8800 | 1.0 | 0.0010 | 0.9999 |
| IBM | 20 | yes | 10,736 | 64 | 10,107 (94.1) | 19 | 565 | 1E-13 | 10 | 4246 | 0.4 | 0.0010 | 0.9974 |
| F<sub>2</sub> | 70 | no | 4,032 | 205 | 3,334 (82.7) | 9 | 493 | 1E-27 | 10 | 4819 | 1.4 | 0.0050 | 0.9996 |
| F<sub>2</sub> | 70 | yes | 4,032 | 301 | 3,336 (82.7) | 11 | 395 | 1E-25 | 10 | 5736 | 1.7 | 0.0050 | 0.9996 |

<sup>6</sup> P-value is the set threshold for grouping markers on the same linkage group. See Methods for details.

**Supplementary Table 8.**
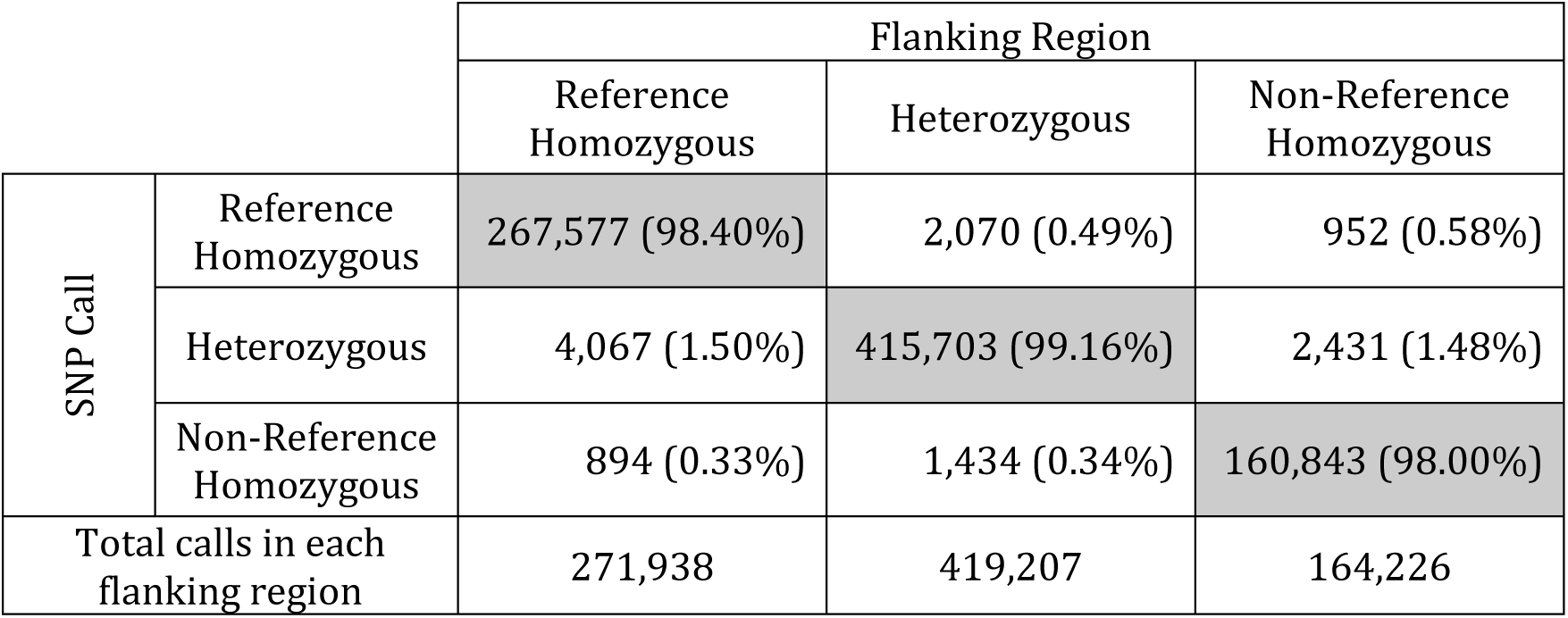
Categorization of tGBS SNP calls by the genotype of the flanking region summed across F_2_ individuals. Accurate calls where the SNP call and the flanking region are in agreement are shaded in gray.

|  |  | Flanking Region |  |  |
| --- | --- | --- | --- | --- |
|  |  | Reference Homozygous | Heterozygous | Non-Reference Homozygous |
| SNP Call | Reference Homozygous | 267,577 (98.40%) | 2,070 (0.49%) | 952 (0.58%) |
|  | Heterozygous | 4,067 (1.50%) | 415,703 (99.16%) | 2,431 (1.48%) |
|  | Non-Reference Homozygous | 894 (0.33%) | 1,434 (0.34%) | 160,843 (98.00%) |
| Total calls in each flanking region |  | 271,938 | 419,207 | 164,226 |

